# Broad-scale signatures of linked selection under divergent recombination landscapes and mating systems in natural populations of rye and barley

**DOI:** 10.1101/2023.02.27.530163

**Authors:** Christina Waesch, Steven Dreissig

## Abstract

Understanding the interactions between patterns of recombination, mating systems, and signatures of natural selection is a central aim in evolutionary biology. Patterns of recombination shape the evolution of genomes by affecting the efficacy of selection. Within populations, genetic shuffling is achieved through meiotic recombination, random chromosome segregation, and the frequency of outbreeding. Recombination landscapes vary between species and populations, and are further influenced by mating systems. Here, we use populations of two related grass species, rye (*Secale cereale*) and barley (*Hordeum vulgare* ssp. *spontaneum*), that differ in their mating system, to analyse signatures of linked selection under divergent recombination landscapes. Rye (outbreeding) and barley (inbreeding) are members of the Poaceae and diverged approximately 15 M years ago. We estimated population recombination rates, analysed patterns of genetic diversity, identified signatures of linked selection, and quantified the fraction and functional class of genes affected by linked selection. In this comparison, we detected signatures of linked selection in low-recombining regions in both species. In inbreeding barley, the low-recombining fraction of the genome was more than 2-fold larger than in outbreeding rye. Furthermore, considering differences in gene density across the genome, approximately 1.5 times more genes were affected by linked selection in barley than in rye. In both species, genomic regions affected by linked selection harbour mostly genes involved in basic cellular processes. We provide empirical evidence for quantitative differences in recombination landscapes between closely related species of divergent mating systems, and discuss the consequences of linked selection and the efficacy of natural selection.

**Significance statement:** The ability of natural selection to disentangle beneficial from deleterious mutations across the genome is shaped by patterns of recombination, yet our understanding of the magnitude of this interaction is incomplete. In our work, we show that patterns of recombination differ between related outbreeding and inbreeding grass species, and that the efficacy of selection differs between them. We provide empirical evidence for quantitative differences in recombination landscapes and patterns of selection.

## Introduction

Linked selection, the effect of positive or purifying selection on genetic variation at linked sites, plays an important role in shaping the evolution of eukaryotic genomes. There are different forms of linked selection depending on the direction of selection. Under the hitchhiking model, a beneficial mutation is rapidly swept to fixation, which leads to a reduction of genetic diversity at linked sites, which are fixed along with the beneficial mutation (Smith & Haigh 2007; Wiehe & Stephan 1993; Slotte 2014). Under the background selection model, purifying selection against deleterious variants leads to a local reduction in genetic diversity at linked sites (Charlesworth et al. 1993; Hudson & Kaplan 1995). In both cases, the underlying recombination landscape, i.e. the frequency distribution of recombination events along the chromosome, plays an important role in shaping the magnitude of linked selection. This leads to the expectation that recombination rates and patterns of nucleotide diversity (*π*) are positively correlated, which was empirically shown in various species (Cutter & Payseur 2013). Signatures of linked selection can be inferred from levels of population differentiation (*F*_*ST*_), since elevated levels of differentiation can arise from both positive and purifying selection and their effects of linked neutral sites (Charlesworth et al. 1997; Keinan & Reich 2010; Muir et al. 2011; Slotte 2014). However, elevated population differentiation could also arise from divergent selection with gene flow between populations (Malinsky et al. 2015; Guerrero & Hahn 2017). In that case, absolute sequence divergence (*d*_*xy*_) between populations is expected to positively correlate with levels of differentiation (*F*_*ST*_). In contrast, linked selection is expected to generate a negative correlation between *F*_*ST*_ and *d*_*xy*_ (Cruickshank & Hahn 2014; Wolf & Ellegren 2017). In recent studies, genomic scans of population differentiation (*F*_*ST*_) were used to identify speciation islands or to characterize patterns of linked selection (Liang et al. 2022; Wang et al. 2022; Shang et al. 2023; Burri et al. 2015; Chase et al. 2021), essentially confirming theoretical predictions regarding the relationship between recombination and patterns of selection (Charlesworth et al. 1997). However, as signatures of selection may vary between populations and species, so may underlying recombination landscapes. Given the increasing availability of large-scale population genetic data sets, chromosome-scale reference genomes, and recombination maps, it is now possible to characterize relationships between recombination and selection in greater detail (Marks et al. 2021; Sun et al. 2022).

Meiosis is a specialized cell division fundamental to eukaryotic reproduction. Patterns of recombination vary along the chromosome, between sexes, individuals, populations, and species (Stapley et al. 2017). Recombination landscapes were shown to be influenced by the spatio-temporal dynamics of meiosis, the local chromatin environment and chromosome structure, recombination rate modifiers, and environmental factors (Henderson & Bomblies 2021; Zelkowski et al. 2019; Brazier & Glémin 2022). In many eukaryotes, recombination rates decrease with distance from the telomeres, and are drastically reduced near centromeres (Haenel et al. 2018; Brazier & Glémin 2022). In most flowering plants, the diversity of recombination landscapes can be explained by the telomere-to-centromere gradient of recombination and the obligatory occurrence of at least one crossover per chromosome (Brazier & Glémin 2022). In a model proposed by Brazier and Glémin (2022), the position of the centromere relative to the telomeres plays and important role in shaping the symmetry of a recombination landscape. In addition to meiotic recombination landscapes, mating systems play an important role in shaping effective recombination rates and patterns of genetic diversity in populations (Jain 1976; Nordborg 2000; Charlesworth & Wright 2001). Based on this, the efficacy of selection is expected to be higher in outbreeding species compared to their inbreeding relatives, under the assumption of otherwise similar recombination landscapes (Charlesworth et al. 1997).

Here, our aim was to characterize signatures of linked selection under divergent recombination landscapes and mating systems. We hypothesized that the interplay between mating system and recombination landscape would have a major impact on the efficacy of selection. This was based on theoretical predictions and empircal observations on the effect of either the local recombination landscape, or the mating system, on signatures of linked selection (Charlesworth et al. 1997; Charlesworth & Wright 2001; Burri et al. 2015; Wang et al. 2022). To adress this question, we used populations of two closely related grass species of divergent mating system, rye (*Secale cereale* L.), and barley (*Hordeum vulgare* L.). Both species divergend approximately 15 M years ago, and rye retained the ancestral outbreeding system, whereas barley evolved into an inbreeding species (Li et al. 2021; Brown et al. 1978; Parzies et al. 2000; Abdel-Ghani et al. 2004; Morrell et al. 2005; Schreiber et al. 2021). We used genotyping-by-sequencing (GBS) data to estimate population recombination rates (*ρ*), analysed patterns of genetic diversity (*π, d*_*xy*_), and identified signatures of linked selection. We detected a two-fold difference in the size of the low-recombining fraction in both species, and approximately 1.5-fold differences in the number of genes affected by linked selection. By analysing gene ontologies of genes in genomic windows under linked selection, we found that, in both species, mostly genes involved in basic cellular functions were are affected by linked selection.

## Results and Discussion

### Recombination landscape variation between inbreeding and outbreeding populations of two closely related species

Within populations, genetic shuffling is achieved through the combined effects of meiotic recombination, random assortment of chromosomes into gametes, and the frequency of outbreeding (Veller et al. 2019; Wang et al. 2019). As such, population-level recombination landscapes, i.e. the chromosomal distribution of meiotic and historical recombination events, are expected to vary between inbreeding and outbreeding species. Rye (genus *Secale*) and barley (genus *Hordeum*) are both members of the Poaceae and diverged approximately 15 M years ago (Li et al. 2021). Whereas wild barley (*Hordeum vulgare* ssp. *spontaneum*) is an inbreeding species, rye retained the ancestral outbreeding mating system (Schreiber et al. 2021; Brown et al. 1978; Parzies et al. 2000; Abdel-Ghani et al. 2004; Morrell et al. 2005).

To explore recombination landscape variation between outbreeding and inbreeding populations of two related plant species, we used genotyping-by-sequencing (GBS) data of 158 individuals of weedy rye (*S. cereale*, 23,628 SNPs, 3.81 SNPs/Mb), and of 114 individuals of wild barley (*H. vulgare* ssp. *spontaneum*, 12,904 SNPs, 2.83 SNPs/Mb) (Milner et al. 2018; Schreiber et al. 2022). We first tested for a comparable distribution of SNPs in both species by counting SNPs in relative chromosomal intervals of 5 %, and observed a correlation of *ρ* = 0.77 between both species (Spearman’s rank correlation, *P* < 2.2 * 10^−16^, Supplementary Figure 1). We then estimated genome-wide effective recombination rates (*ρ/kb*), which revealed substantial variation with a genome-wide average of 14.32 *ρ/kb* in weedy rye and 0.03 *ρ/kb* in wild barley (*P* = 6.7*10^−98^, Wilcoxon-Mann-Whitney *U*-test, Figure 1A). Since measures of population-level recombination rates are influenced by the underlying mating system, we estimated individual inbreeding coefficients (*F*) in both species, and observed an average of 0.02 in weedy rye and an average of 0.98 in wild barley, reflecting their mating systems and differences in recombination rates (*P* = 3.7*10^−44^, Wilcoxon-Mann-Whitney *U*-test, Figure 1B). In weedy rye, genome-wide nucleotide diversity was found significantly higher than in wild barley (*π*_*rye*_ = 1.18 * 10^−6^; *π*_*barley*_ = 7.57 * 10^−7^; *P* = 2.7*10^−35^, Wilcoxon-Mann-Whitney *U*-test, Figure 1C).

**Figure 1.**
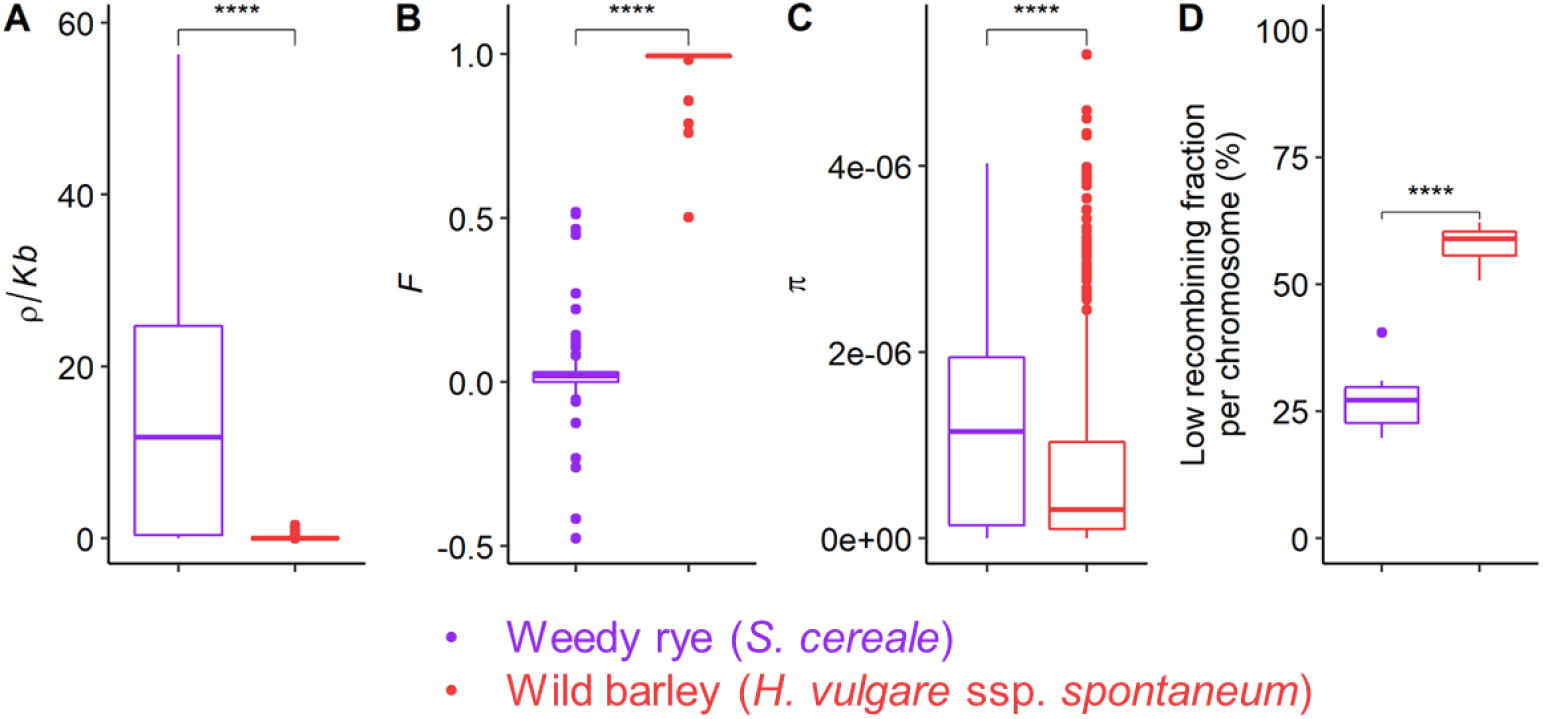
Estimates of effective recombination rate, inbreeding, and nucleotide diversity in weedy rye and wild barley. As a self-incompatible, outbreeding species, weedy rye showed higher population recombination rates (*ρ/kb*), reduced inbreeding (*F*), higher nucleotide diversity (*π*), but also a smaller low-recombining fraction per chromosome. (A) population recombination rate (*ρ/kb*) in 10 Mb windows. (B) Individual inbreeding coefficient (*F*). (C) Nucleotide diversity (*π*) in overlapping windows of 10 Mb, step size of 2.5 Mb. (D) Low-recombining fraction per chromosome, defined as 10 Mb windows with 20-fold reduced *ρ/kb* compared to chromosome-wide mean. ***: *P* <= 0.001; Wilcoxon-Mann-Whitney *U*-test).

Next, we asked whether recombination landscapes, i.e. the distribution of low- and high-recombining regions along the chromosome, differed between weedy rye and wild barley. Given a mean SNP density of 3.81 (rye) and 2.82 (barley) SNPs/Mb in our GBS data, we aggregated mean *ρ/kb* values in non-overlapping windows of 10 Mb, which resulted in at least 1 SNP per window, even across peri-centromeric regions. Low-recombining windows were defined by a 20-fold reduction compared to the respective chromosome-wide average (Choo 1998; Baker et al. 2014; Schreiber et al. 2022). This revealed substantial differences in the fraction of the genome that is low-recombining, with the low-recombining fraction amounting to 27.44 % of the genome in weedy rye and 57.66 % in wild barley (Figure 1D).

LD-based estimates of population recombination rates are influenced by demographic history, where population bottlenecks (e.g. domestication events) would cause lower effective recombination rates caused by a reduction in the effective population size (*Ne*) (Torres et al. 2020; Beissinger et al. 2016). On the other hand, broad-scale LD-based estimates were shown to correlate well with per-generation meiotic recombination rates (cM/Mb) in plants estimated based on linkage maps (Choi et al. 2013; Dreissig et al. 2019; Marand et al. 2019; Danguy des Déserts et al. 2021; Fuentes et al. 2021). Based on this, we focused on weedy and wild populations that did not undergo recent demographic changes such as domestication bottlenecks (Sun et al. 2021; Badr et al. 2000; Kilian et al. 2006; Russell et al. 2016). Recombination landscapes were shown to vary within and between plant and animal species, with rank correlations ranging from 0.4 to > 0.9 between populations within species (Bauer et al. 2013; Dreissig et al. 2019; Spence & Song 2019; Schreiber et al. 2022; Danguy des Déserts et al. 2021). In previous work, we characterized intraspecific recombination landscape variation among populations of rye, and found that the fraction of low-recombining regions varied from 24.2 % to 42.1 % between weedy and domesticated rye (Schreiber et al. 2022). Many comparisons between related species were focused on crossover frequency or genome-wide recombination rate (reviewed by (Stapley et al. 2017; Mercier et al. 2015; Ross-Ibarra 2004)), and recent studies were focused on the distribution of recombination events along the chromosome (Brand et al. 2018; Cai et al. 2023; Brazier & Glémin 2022; Haenel et al. 2018). According to a recent meta-study, recombination landscapes in flowering plants are best explained by distance to telomere and centromere effects, as well as one obligatory crossover per chromosome (Brazier & Glémin 2022). Both rye and barley chromosomes are metacentric, submetacentric, or acrocentric, and rye chromosome are 100 – 200 Mb longer than barley chromosomes (Künzel et al. 2000; Bennett et al. 1977; Mascher et al. 2017; Rabanus-Wallace et al. 2021). Another model proposed by Haenel et al. (2018), which was derived from analysing recombination landscapes in 62 animal, plant, and fungal species, stated chromosome length as a major predictor of recombination landscapes, with larger chromosomes having larger low-recombining regions. Our observations do not strictly adhere to the chromosome-length model proposed by Haenel et al. (2018), but are rather in agreement with the telomere-centromere-position model of Brazier and Glémin (2022), where variations in telomere to centromere distance have an effect on the size of low-recombining regions and the symmetry of recombination landscapes.

### Recombination landscape variation shapes patterns of linked selection

To analyse signatures of linked selection, owing either to background selection or hitchhiking, we selected an outgroup of approximately equal size for each species and estimated absolute sequence divergence (*d*_*xy*_) and genetic differentiation (*Z(F*_*ST*_)) between subpopulations. In case of rye, we selected 120 wild individuals of *S. strictum*. That taxon is a perennial outbreeder closely related to *S. cereale*, genetically distinct from *S. cereale*, and commonly recognized as a separate species (Figure 2A) (Schreiber et al. 2019; Sun et al. 2021; Schreiber et al. 2021). For wild barley, we selected another 117 individuals from a geographic subcluster in the eastern Fertile Crescent (Figure 2B). In both species, overall genetic differentiation between pairwise subpopulations was similar, with a genome-wide average *F*_*ST*_-value of 0.269 between weedy rye and wild *S. strictum*, and 0.312 between wild barley western and eastern subpopulations.

**Figure 2.**
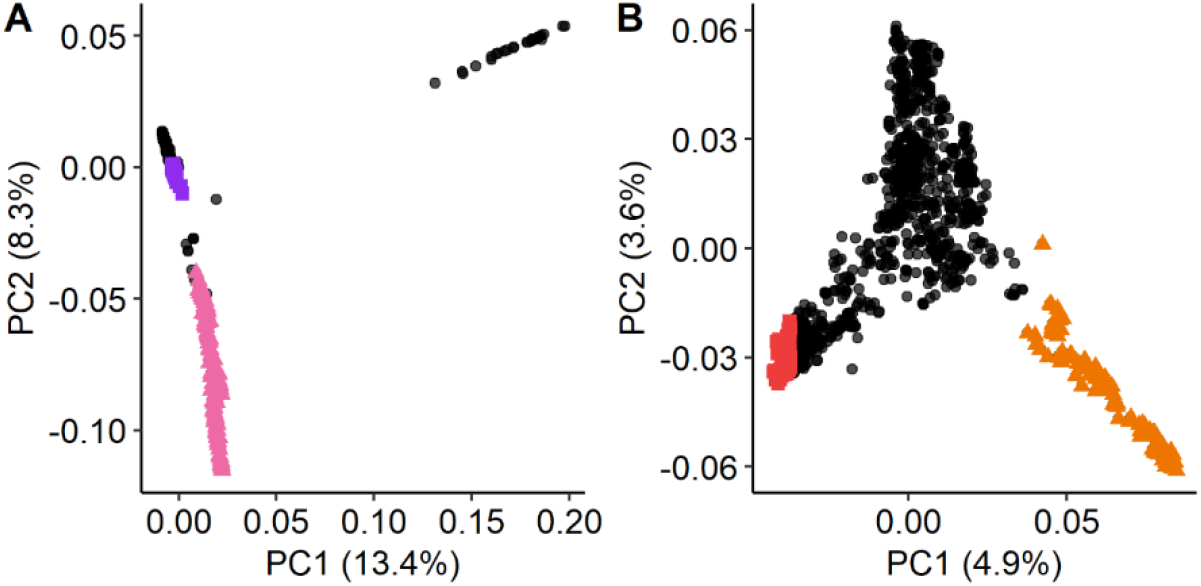
Population structure and selection of outgroups in rye and barley. (A) Principal component analysis showing relationships among genetic clusters of rye. Weedy rye (*S. cereale*) individuals are shown in purple, whereas wild *S. strictum* individuals are shown in pink (based on (Rabanus-Wallace et al. 2021). (B) PCA showing structure among wild barley accessions. The focal subpopulation is shown in red (western), whereas another subpopulation, which was used as an outgroup, is shown in orange (eastern) (based on (Milner et al. 2018)).

Next, we estimated genetic differentiation (*Z(F*_*ST*_)), absolute sequence divergence (*d*_*xy*_) between subpopulations, and within-population nucleotide diversity (*π*) in overlapping windows of 10 Mb (2.5 Mb step size), which revealed that nucleotide diversity and sequence divergence were reduced, and genetic differentiation increased, in low-recombining regions (Figure 3A, 3B). In both species, we observed negative correlations between genetic differentiation and recombination rates (Figure 3C, 3D; Spearman’s *ρ*_*rye*_ = -0.52; Spearman’s *ρ*_*barley*_ = -0.51; *P* < 2.2 * 10^−16^), and positive correlations between sequence divergence and recombination rates (Figure 3C, 3D; Spearman’s *ρ*_*rye*_ = 0.9; Spearman’s *ρ*_*barley*_ = 0.73; *P* < 2.2 * 10^−16^).

**Figure 3.**
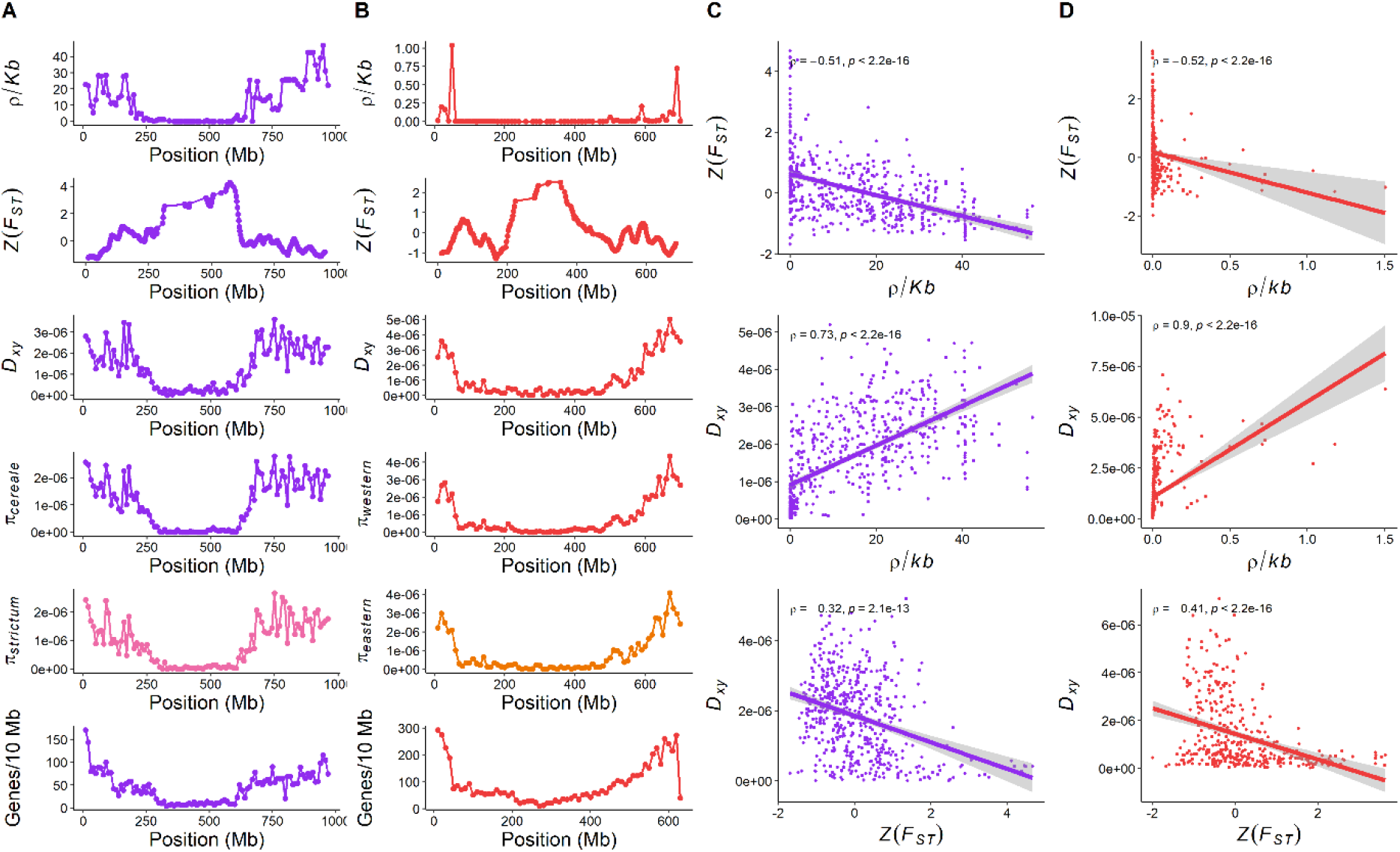
Broad-scale patterns of linked selection in weedy rye and wild barley. Patterns of effective recombination rates (*ρ/kb*), genetic differentiation (*Z(FST)*), absolute sequence divergence (*d*_*xy*_), nucleotide diversity (*π*), gene density shown for chromosome 3R/3H in weedy rye (A, C, purple) and wild barley (B, D, red) reveal broad-scale signatures of linked selection in low-recombining peri-centromeric regions. All genomic features were aggregated in overlapping windows of 10 Mb with a step size of 2.5 Mb. *ρ* = Spearman’s rank correlation coefficient.

In low-recombining regions, patterns of genetic differentiation are expected to arise through linked selection processes, either background selection or hitchhiking, where positive or negative selection leads to a reduction of diversity at linked neutral sites (Charlesworth et al. 1997; Cutter & Payseur 2013; Slotte 2014). Patterns of genetic differentiation, however, could also arise through divergent selection and gene flow between subpopulations. In that case, a positive correlation between sequence divergence and genetic differentiation is expected (Cruickshank & Hahn 2014; Burri et al. 2015). In our analyses, we observed negative correlations between sequence divergence and genetic differentiation in both species, suggesting that both species genome’s were shaped by linked selection (Figure 3C, 3D; Spearman’s *ρ*_*rye*_ = -0.41; Spearman’s *ρ*_*barley*_ = -0.32; *P* < 2.2 * 10^−16^).

### Distribution of gene density in low-recombining regions differs between outbreeding rye and inbreeding barley

Under divergent recombination landscapes and effective recombination rates, the question arises as to how many genes are located in low-recombining regions, which might in turn be affected by linked selection. In order to address this question, we aggregated the number of annotated high-confidence genes in non-overlapping 10 Mb windows in both species. In weedy rye, 9.7 % of all genes were located in low-recombining regions, whereas 30.2 % of genes were located in low-recombining regions in wild barley (Figure 4A). Given that the fraction of the genome which is low-recombining differed between both species, we calculated the ratio of the proportion of genes in low-recombining regions to the low-recombining fraction per chromosome. If genes were distributed evenly along the chromosome, it would result in a ratio of 1. In that case, the fraction of genes potentially affected by linked selection would be influenced by the size of the low-recombining fraction alone. This analysis revealed a ratio of 0.35 in weedy rye and a ratio of 0.52 in wild barley, showing that in wild barley, a larger fraction of genes is potentially affected by linked selection (Figure 4B).

**Figure 4.**
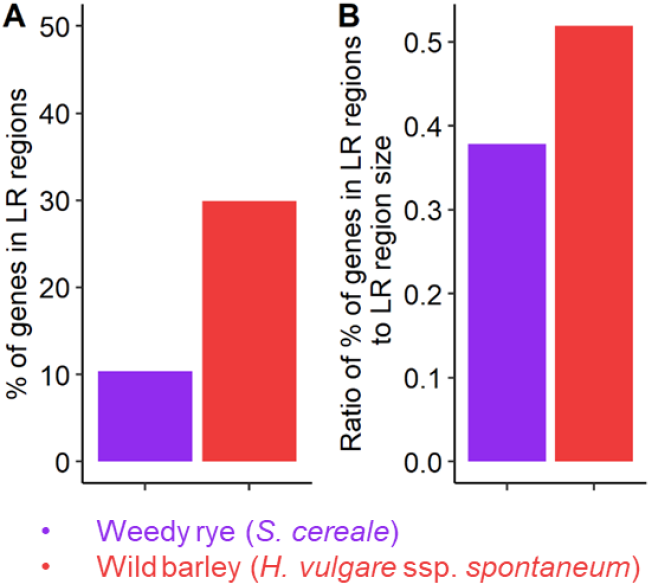
Gene density in low-recombining regions. **(A)** Percentage of genes located in low-recombining windows in weedy rye and wild barley. **(B)** Ratio of the percentage of genes in low-recombining regions to the size of the low-recombining fraction.

However, a difference in the fraction of genes affected by linked selection alone does not explain whether or not the efficacy of natural selection is hampered by larger low-recombining regions, as it depends on which genes are located within and whether these are the targets of natural selection. In order to address this question, we investigated which functional classes of genes were affected by linked selection by performing gene ontology (GO) enrichment analysis for genes extracted from genomic windows showing strong signatures of linked selection (window size = 10 Mb, *Z(F*_*ST*_) scores => 0.95 percentile).

We identified a total of 29 and 23 genomic windows harbouring an average of 26.9 and 31.9 genes per window in rye and barley, respectively. In total, this amounted to 780 rye genes and 733 barley genes underlying *F*_*ST*_ peaks. Our GO enrichment analysis revealed that 143 and 70 GO-terms were significantly (p < 0.05) enriched in regions affected by linked selection in rye and barley, respectively. Among these, most were involved in basic cellular processes, such as lysine metabolism, glucose metabolism, photosynthesis, translation, as well as actin and ribosome function (Figure 5). This might be expected, at least in barley, since it was shown that peri-centromeric, low-recombining regions were enriched in genes related to photosynthesis and translation (Mascher et al. 2017). However, we cannot confirm whether or not these genes are the actual targets of natural selection. Nevertheless, our data suggest that many genes involved in basic cellular functions are potentially affected by linked selection due to their genomic location. Interestingly, a similar pattern is known in pathogenic fungi, known as the ‘two-speed genome’ concept, where genes in gene sparse, repeat rich regions show faster rates of adaptive evolution (Dong et al. 2015; Raffaele & Kamoun 2012; Croll & McDonald 2012). In future, high-coverage whole-genome-sequencing, preferably in combination with long-read sequencing to detect structural variations, might enable us to identify the actual targets of natural selection in these species.

**Figure 5.**
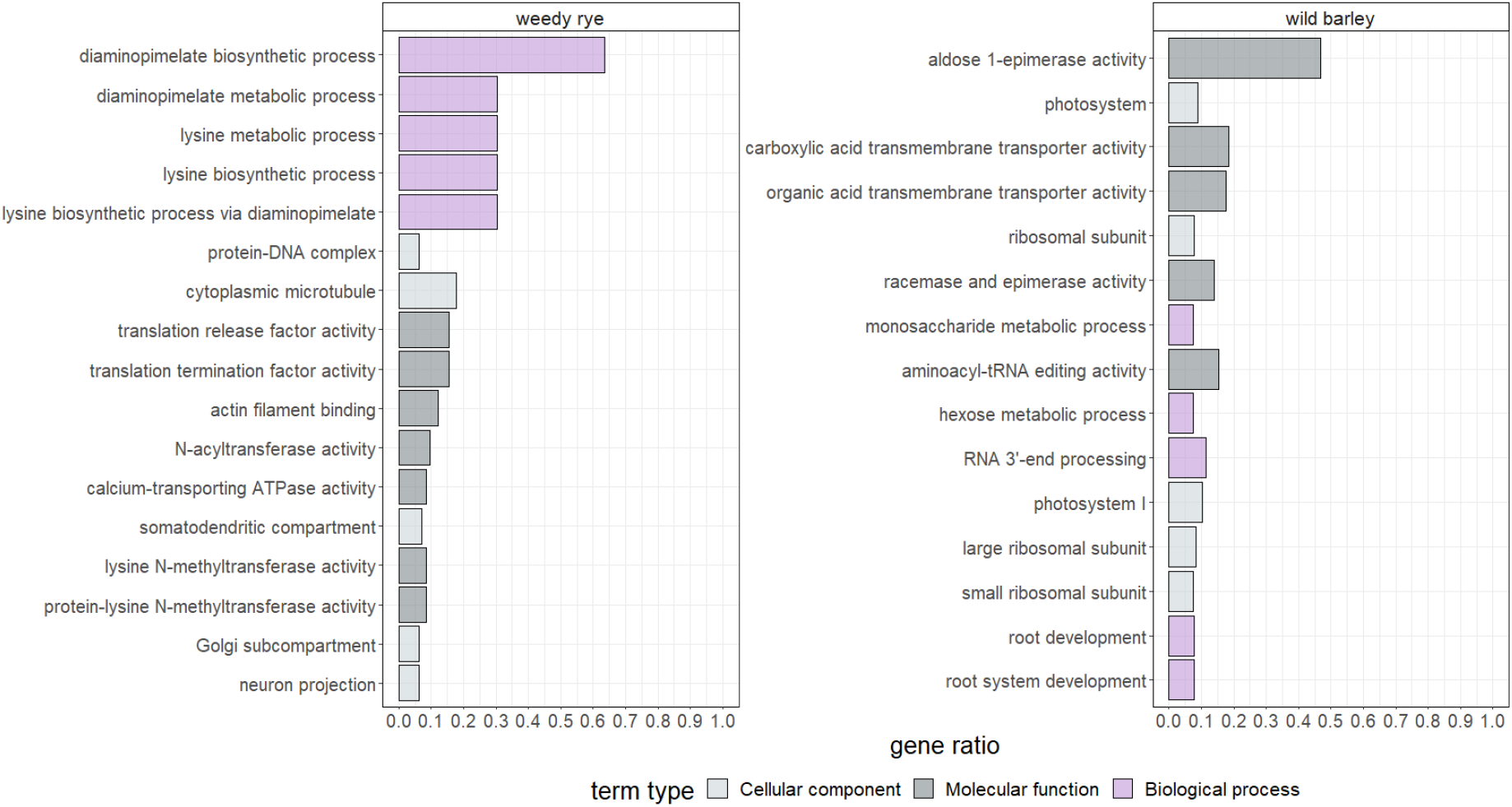
Gene ontology enrichment in genomic regions affected by linked selection. In both species, regions affected by linked selection are enriched in genes involved in basic cellular processes. Significantly enriched GO-terms are shown for cellular components (light grey), molecular functions (dark grey), and biological processes (purple) and sorted by *P*-value. Gene enrichment ratio was calculated as the ratio of GO-terms in high *F*_*ST*_ regions to the total occurrence in the reference genome / vs the amount of high-confidence genes in the entire genome. Only the top five GO-terms for gene ratio of each category are shown.

Taken together, our work showed broad-scale signatures of linked selection in natural populations of outbreeding rye and inbreeding barley, evident as negative correlations between recombination rates (*ρ/Kb*) and genetic differentiation (*F*_*ST*_), positive correlations between absolute sequence divergence (*d*_*xy*_) and *ρ/Kb*, and negative correlations between *d*_*xy*_ and *F*_*ST*_. Furthermore, we revealed substantial differences in genome-wide effective recombination rates as well as recombination landscapes, expressed as the low-recombining fraction of the genome. Given that, in wild barley, comparatively more genes were located in low-recombining regions (1.5 times more), effective recombination rates were drastically lower (*ρ/Kb* = 0.03 vs. *ρ/Kb* = 14.32), and low-recombining regions larger (2 times larger), we provide empirical evidence for quantitative differences in the efficacy of selection in two closely related grass species.

**Supplementary Figure 1.**
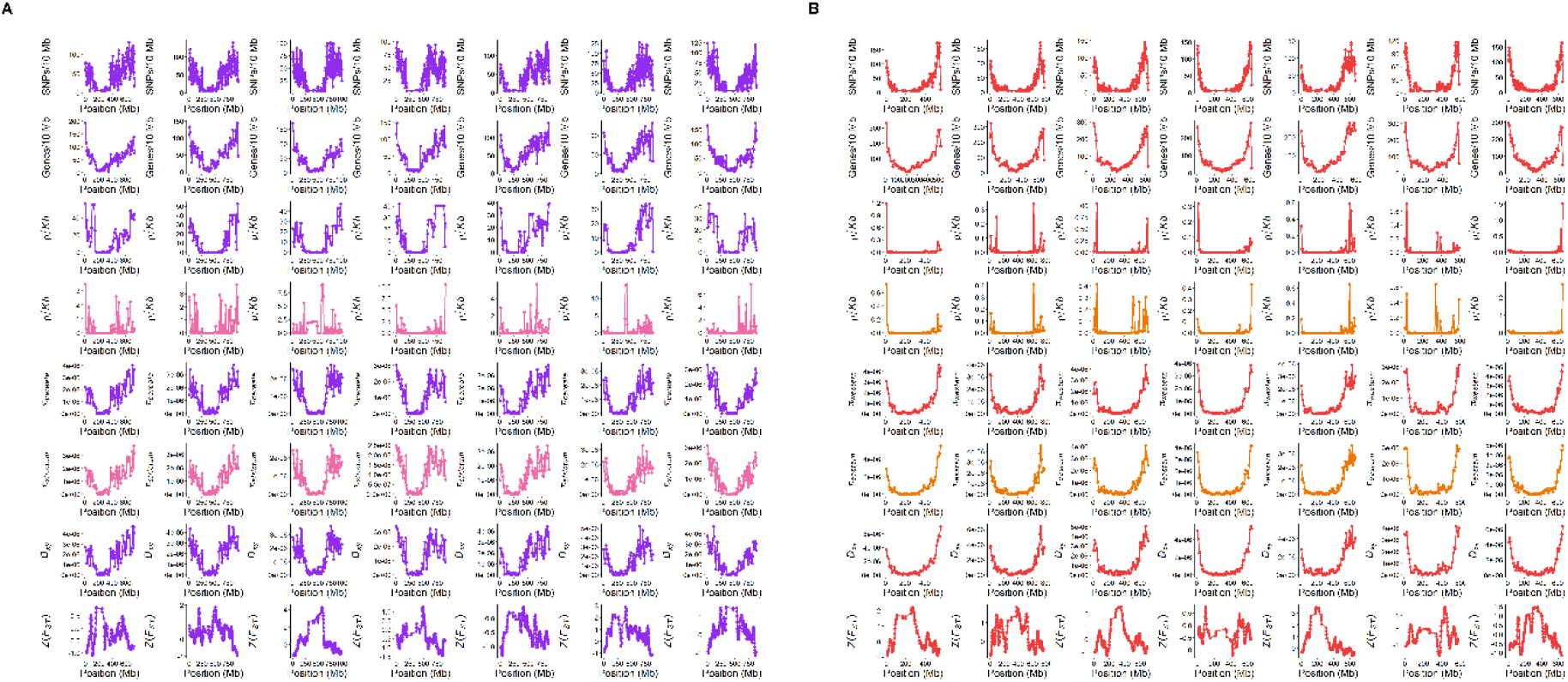
Summary statistics in rye (A) and barley B).

## Materials and Methods

### Plant material, genome sequence data, and population structure analysis

We used genotyping-by-sequencing (GBS) data of two large diversity panels of rye (genus *Secale*) and wild barley (*H. vulgare* ssp. *spontaneum*) comprising 1397 and 1140 accessions, respectively (Milner et al. 2018; Schreiber et al. 2022). Rye GBS and passport data were retrieved from the European Variant Archive under project number PRJEB5152, and barley GBS and passport were retrieved from e!DAL (https://doi.org/10.5447/IPK/2018/9). Original variant matrices comprised 58,688 and 171,263 polymorphic sites for rye and barley, respectively. Both matrices were filtered for minor allele frequencies > 0.1, maximum missing data < 0.1, to contain only biallelic sites, and no insertions/deletions using VCFtools (Danecek et al. 2011). Filtered SNP matrices were used to perform principal component analyses using SNPrelate (algorithm=“exact”, eigen.method=“DSPEVX”) (Zheng et al. 2012). Based on passport data and PCA results, we separated the rye panel into two subpopulations, one comprising 65 weedy rye (*S. cereale*) accessions (total of 158 individuals, mean of 2.4 individuals per accession) and an outgroup comprising 21 *S. strictum* accessions (total of 120 individuals, mean of 5.7 individuals per accession). The wild barley diversity panel was separated into two subpopulations based on passport data and PCA results, with one subpopulation from the south-western part of the Fertile Crescent comprising 114 accessions (1 individual per accession) (hereafter named western), and another subpopulation from the south-eastern part of the Fertile Crescent comprising 117 accessions (1 individual per accessions) (hereafter named eastern). Final SNP matrices of selected accessions contained 23,628 polymorphic sites in rye subpopulations, and 12,904 polymorphic sites in wild barley subpopulations. A list of all individuals included in the analysis is provided in Supplementary File 1.

### Recombination landscape analysis

We used single-nucleotide-polymorphism (SNP) data to estimate population recombination rates (ρ/Kb, ρ = 4N_e_ * r) based on coalescent theory via LDhat (Stumpf & McVean 2003; Auton & McVean 2007). LDhat was run for 10,000,000 iterations, a block penalty of 5, a population scaled mutation rate (*θ*) of 0.01, sampling every 5000 runs, and the first 100,000 runs were discarded as burn-in. ρ/Kb values were aggregated in non-overlapping genomic windows of 10 Mb along the chromosome. Low-recombining regions were defined as 10 Mb windows showing 20-fold lower values than the respective chromosome-wide mean (Choo 1998; Baker et al. 2014; Fuentes et al. 2021; Schreiber et al. 2022). Recombination rate estimates are provided in Supplementary File 1.

### Gene density, nucleotide diversity (*π*), absolute sequence divergence (*d*_*xy*_), and genetic differentiation (*F*_*ST*_)

Gene density and all diversity statistics were aggregated in overlapping genomic windows of 10 Mb with a step size of 2.5 Mb. We calculated gene density based on high-confidence gene models in the barley reference genome assembly of the cultivar ‘Morex’ v2 (Mascher et al. 2017) and the rye ‘Lo7’ reference genome assembly v1 (Rabanus-Wallace et al. 2021). Nucleotide diversity (*π*), absolute sequence divergence (*d*_*xy*_), and genetic differentiation (*F*_*ST*_) were calculate using the PopGenome package in R (Pfeifer et al. 2014). *F*_*ST*_-values were Z-transformed using the following formula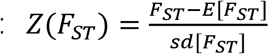 where *E*[*F*_*ST*_ ]and *sd*[*F*_*ST*_ ]denote the expected value and standard deviation of *F*_*ST*_.

### Gene ontology enrichment analysis

Gene ontology (GO) enrichment analysis was done using agriGO v.2.0 (Tian et al. 2017) via singular enrichment analysis (SEA). For both species, we extracted gene IDs and GO-terms of genes located in 10 Mb windows showing *Z(F*_*ST*_)-values above the chromosome-specific 0.95 percentile. Singular enrichment analysis was performed using gene IDs and GO-terms of all high-confidence genes as reference (34,433 high-confidence genes in rye, 32,782 high-confidence genes in barley). Statistical tests were conducted using Fisher’s exact test and multiple-test adjustment method after Yekutieli (significance level of 0.05), with minimum number of entries set to 5. Gene enrichment ratios were calculated as the ratio of significant (p < 0.05) enriched genes within the 10 Mb window of elevated *F*_*ST*_-values against the amount of high-confidence genes in the entire genome. Gene IDs and GO-terms are provided in Supplementary File 1.

## Data availability

Rye and barley accessions used in this study are listed in Supplementary File 1.

## Author contributions

Conceptualization: S.D., Methodology: C.W. and S.D., Software: C.W. and S.D., Validation: C.W. and S.D., Formal Analysis: C.W. and S.D., Investigation: C.W. and S.D., Resources: S.D., Data Curation: S.D., Writing – Original Draft: C.W. and S.D., Writing – Review & Editing: C.W. and S.D., Visualization: C.W. and S.D., Supervision: S.D., Project Administration: S.D., Funding Acquisition: S.D.

## Acknowledgements

We would like to thank Martin Mascher and Markus Stetter for insightful discussions. We are also grateful to all authors who made genome sequence data of barley and rye publicly available.

